# Superadditive and subadditive dynamics are not inherent to the types of interacting threat

**DOI:** 10.1101/522755

**Authors:** V Haller-Bull, M Bode

## Abstract

Species and ecosystems usually face more than one threat. The damage caused by these multiple threats can accumulate nonlinearly: either subadditively, when the joint damage of combined threats is less than the damages of both threats individually added together, or superadditively, when the joint damage is greater than the two individual damages added together. These additivity dynamics are commonly attributed to the nature of the threatening processes, but conflicting empirical observations challenge this assumption. Here, we use a theoretical model to demonstrate that the additivity of threats can change with different magnitudes of threat impacts (effect of a threat on the population parameter, like growth rate). We use a harvested single-species population model to integrate the effects of multiple threats on equilibrium abundance. Our results reveal that threats do not always display consistent additive behavior, even in simple systems. Instead, their additivity depends on the magnitudes of the impacts of two threats, and the population parameter that is impacted by each threat. In our model specifically, when multiple threats have a low impact on the growth rate of a population, they display superadditive dynamics. In contrast, threats that impact the species’ carrying capacity are always additive or subadditive. These dynamics can be understood by reference to the curvature of the relationship between a given population parameter (e.g., growth) and equilibrium population size. Our results suggest that management actions can achieve amplified benefits if they target low-amplitude threats that affect the growth rate, since these will be in a superadditive phase. More generally, our results suggest that cumulative impact theory should focus more than previously on the magnitude of the impact on the population parameter, and should be cautious about attributing additive dynamics to particular threat combinations.

## Introduction

Species and ecosystems across the globe are exposed to a large variety of threats. Coral reefs, for example, face a variety of threats, including direct marine threats such as fishing; land-based threats such as water quality; and global threats such as coral bleaching (1, 2). When threats occur in conjunction, the total damages caused can interact and display nonlinear behaviors (3), where the presence of a given threat can be magnified, reduced, or erased by the presence of another threat. These cumulative threat dynamics have important implications for the benefits of management actions (2, 4, 5).

Since we cannot investigate all moving components simultaneously, we narrow down the issue to a single population faced by several threats. When a threat on a population occurs it passes through several stages before we see the damage (Fig.1). In this paper, we distinguish between the terms “threat”, “impact” and “damage”. We use the term threat categorically, to refer to the nature of the process that affects our population, such as fishing, or eutrophication. The impact is defined as the actual reduction of a population parameter that the threat causes (Table 1). The damage is defined as the change in the population size that results from that impact. For example, we could have a cyclone (a threat) occurring at a reef. This cyclone might reduce the amount of habitat available for the fish population, i.e. the carrying capacity is reduced. This reduction of the fishes’ carrying capacity is the actual impact on the population. Damage on the other hand is the effect of the cyclone that we can measure at some point after the threat has occurred, usually this is a population reduction. While these two concepts are often used interchangeably, they need to be distinguished to avoid confusion and also mismatches between the results of different studies. As this paper will show additive impacts does not always lead to additive damages. This can especially be an issue when considering conservation since management goals are usually based on population sizes (i.e. damages are of interest) and not population internal parameters (i.e. impacts).

**Figure 1.**
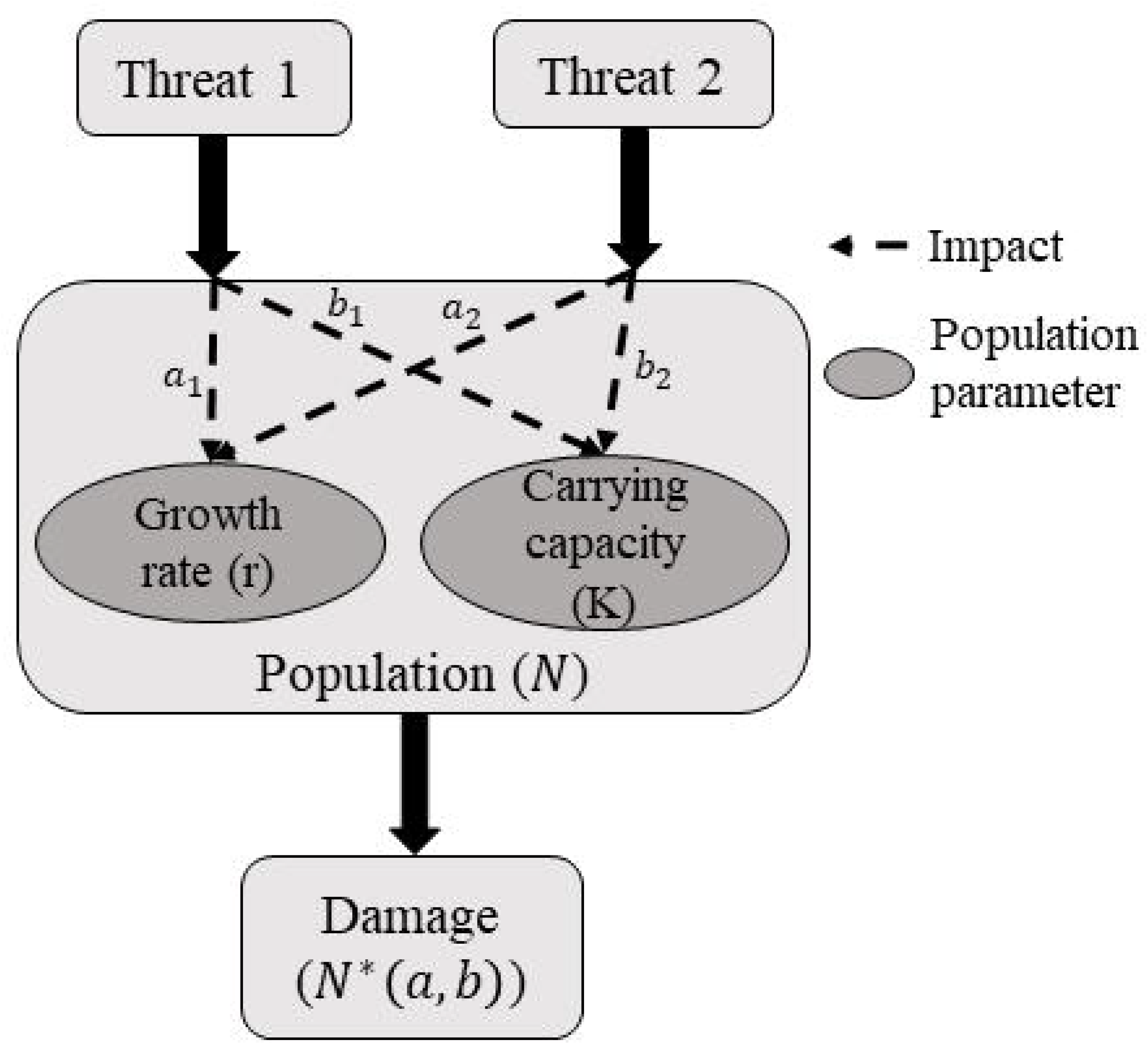
Schematic diagram of the stages through which threats impact populations.

**Table 1.**
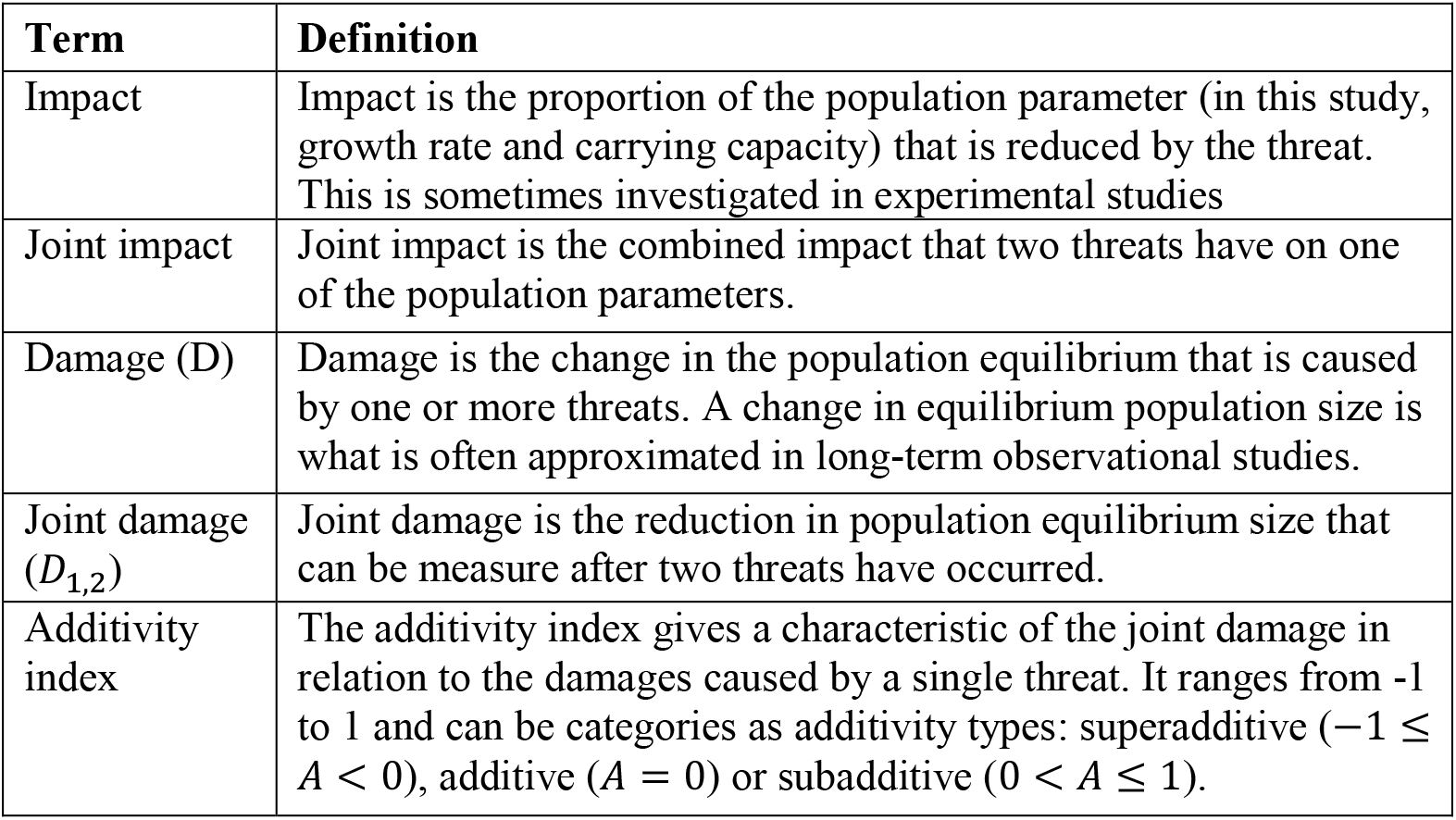
Definition and connection of commonly used terms in this paper.

| Term | Definition |
| --- | --- |
| Impact | Impact is the proportion of the population parameter (in this study, growth rate and carrying capacity) that is reduced by the threat. This is sometimes investigated in experimental studies |
| Joint impact | Joint impact is the combined impact that two threats have on one of the population parameters. |
| Damage (D) | Damage is the change in the population equilibrium that is caused by one or more threats. A change in equilibrium population size is what is often approximated in long-term observational studies. |
| Joint damage ($D_{1,2}$) | Joint damage is the reduction in population equilibrium size that can be measure after two threats have occurred. |
| Additivity index | The additivity index gives a characteristic of the joint damage in relation to the damages caused by a single threat. It ranges from -1 to 1 and can be categories as additivity types: superadditive ($-1 \leq A < 0$), additive ($A = 0$) or subadditive ($0 < A \leq 1$). |

When multiple threats occur simultaneously (6), the damage they cause to a species’ population(or ecosystem feature such as species richness) is a result of each individual threat, and the interaction between them (5, 7). Interactions can occur in a variety of ways; here, we focus solely on interactions that occur within a single population. The accumulated damage of multiple threats are not always additive, meaning two threats occurring simultaneously aren’t always the same as the individual damages added together. The existence of non-additive threat interaction dynamics has been shown repeatedly (7–10). Since threats act onto several stages within a population (impact and damage), non-additive interactions can also occur on both of these stages. This means that we can have both interactions in the impacts or interactions in the damages. Both of these are treated separately in this study.

Some experimental studies measure a process within the population like the growth rate and try to determine whether the damages of two threats on this rate are additive, or non-additive. This is a study of the impacts. However, when we have an observational study that measures the reduction in population size over a certain time period in the presence and absence of different combinations of threats then we are concerned with the non-additivity of the damages. In this paper, we fix the interaction within the threat impact (either additive or multiplicative) and measure the interaction of the damages (additive, subadditive or superadditive). Subadditivity (often called compensatory effects) occurs when the damages caused by the combined threat impacts, D(A&B), are smaller than the sum of the damages of the individual threat impacts, D(A&B) < D(A)+D(B) (11). Similarly, superadditivity (often called synergistic effects) is defined as the joint damage being larger than the sum of the individual damages, D(A&B) > D(A)+D(B) (11).

The majority of studies on threat interactions rely on experimental or observational methods. The main aim of these studies is to identify which threat combination (e.g., deceased salinity and elevated temperature) displays which type of additivity. However, the studies often disagree about the type of additivity, even when considering the same study species and threats. Crain and Kroeker (7) reviewed 202 studies on these interaction types in marine systems and found that 26% of threat combinations create additive damages, while 36% are superadditive and 38% are subadditive. Note here that these studies did not differentiate between impacts and damages. There was also variation within threat combinations: all threat combinations that had been thoroughly investigated displayed all three additivity types (7). For example, 34 independent factorial experiments that investigate the additive behavior of UV light and fishing found additive behavior in 17 cases, subadditive behavior in 5 cases and superadditive behavior in 12 cases (7). So far, this variation has been explained by context dependence (7), including the number of threats considered and the trophic level of the species experiencing the threat. Here, we investigate an alternative explanation for the observed variation: additivity of joint damages of threats can change with varying the magnitude of impacts.

Investigating different magnitudes of threat impacts is difficult in both observational and experimental studies because it would require a large amount of data or a very complicated and extensive experimental design, with the species or community being exposed to the threats individually and in combination, across a range of impact magnitudes. For example, Schlöder and D’Croz (12) investigated the damage ocaused bytemperature and nitrate on two coral species, *Pocillopora damicornis* and *Porites lobate*. In this experiment, 60 coral pieces were grown for 30 days in isolation and the frequency and volume of their zooxanthellae was measured. The magnitude of the threat impacts was only classified in two (nitrate) or three (temperature) categories, resulting in six possible scenarios and leaving five replicates per species. Even an increase to three levels in nitrate would result in an increase of the combinations to nine combinations and 90 coral fragments if keeping the replication constant. This makes the investigation of many different magnitudes of threat impacts very challenging; modelling studies need to be used to address these kind of questions more holistically. Simulations of threats at many trophic levels, including all of their types and magnitudes of impacts, as well as utilising large sample sizes can be analysed to draw more general conclusions. In contrast, models can evaluate threats and their management in situations where manipulation or experimentation is challenging (5, 13, 14).

In this study, we analyse the conditions within a population and their threats that lead to superadditive and subadditive behavior in damages. Our aim is to theoretically investigate the additivity of joint threat impacts, and to offer a more nuanced understanding of the factors that influence additivity. We are especially interested in understanding how additivity varies with different magnitudes of impacts, and how it depends on the parameters impacted by those threats. We use a suite of single-species population models to simulate damages caused by threats in isolation and combination to identify the interaction behavior. Then, we identify and explain the conditions that lead to super- and subadditivity damages. Finally, possible management actions are simulated and their relative benefits depending on the additivity are compared.

## Methods

Our analyses are based on a single-species population model – the harvested logistic model (Eq. 1) – which allows us to derive analytical results, and to more easily interpret them in the context of cumulative threat theory. Impacts are modelled as proportional reductions (*a* and *b*) in two population parameters: the growth rate (*r*), and the carrying capacity (*K*) respectively. So the logistic model

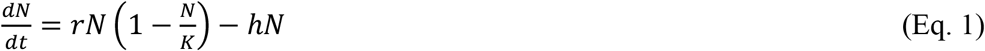

becomes

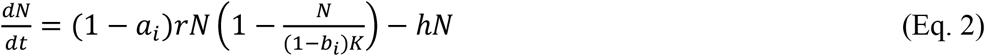

in the presence of threat i.

We note that in this model, a single threat i can impact both our population parameters. For example, in coral reef ecosystems sedimentation simultaneously reduces the habitat available to corals (*K*), and increases coral mortality (*r*). Our approach would therefore allow this single threat to interact with itself. In Eq. 1 and Eq. 2, *r* = growth rate, *a*_*i*_ = impact of threat i on the growth rate, *K* = carrying capacity, *b*_*i*_ = impact of threat i on carrying capacity, *h* = harvest rate, also 0 ≤ *a*_*i*_ ≤ 1, 0 ≤ *b*_*i*_ ≤ 1,0 ≤ *r* ≤ 1,0 ≤ *h* ≤ 1 and *h* < *r*. A value of zero for either *a*_*i*_ or *b*_*i*_therefore indicates no impact of threat i on the parameter, while a value of one indicates a total loss of the process represented by the parameter. Our analyses focus on the equilibrium population in the face of two threats that each impact one or both a population parameters to create a new population equilibrium,

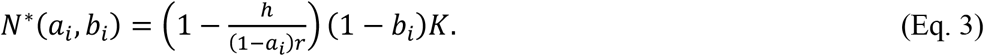

If *N*^∗^(*a*_*i*_, *b*_*i*_) < 0, we consider the population to be extinct and set *N*^∗^(*a*_*i*_, *b*_*i*_) = 0. In the figures a line is added that separates extinct from extant populations.

Furthermore, we define the damage (*D*) to be the reduction in population size caused by the threat:

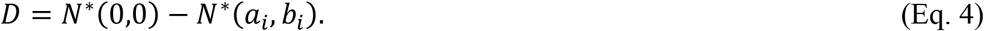

We could model the impact of two threats (later referred to as threat 1 and threat 2) on a single parameter in two ways: multiplicative (1 − *b*) ∗ (1 − *b*) or additive 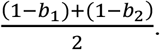 Both of these are reasonable. Additive impacts would indicate that the two impacts occur independently and simultaneously with no influence on one another, for example, a new port being build, removing a certain amount of the coral reef habitat. At the same time an oil tanker spills nearby and more habitat becomes unlivable. Both of these threats (port and oil spill) act independently so impact the population process of carrying capacity without regard for one another. Multiplicative impacts would indicate that they change the impact of one another, for example, if they acted consecutively, the impact of threat 2 therefore affects a parameter that has already been impacted by threat 1. In our previous example this would mean that the oil spill occurs close to (and potentially after) the new port, so that only the habitat remaining after threat 1 (port construction) is impacted by threat 2 (oil spill). Which model is most appropriate depends on the threats and how they affect the surroundings and physiology of the modelled organisms. Here we present the results of the additive model, however the analysis for the multiplicative version can be found in the supplementary materials. Furthermore, the supplementary materials also provide the results for the equivalent analysis for the Beverton-Holt (S2) and the Ricker model (S3).While there are slight differences in the results, all major conclusions in this paper are supported by the results of all analyses.

To categorise the joint damage caused by multiple threats we have created an additivity index (A). It is based on the population equilibria in the presence and absence of the threats. Basically, the additivity index is equal to the sum of the damage caused by each threat and its impacts separately, minus the damage caused by both threats simultaneously,

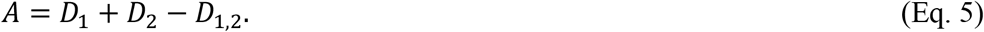

When A is negative the joint damage is superadditive; when A is positive then the joint damage is subadditive.

We consider four different types of interacting threats (Table 2). In our first two cases, both threats impact only one parameter, either the carrying capacity ( *a*_1_ = *a*_2_ = 0, *b*_1_, *b*_2_ ≠ 0) or the growth rate ( *a*_1_, *a*_2_ ≠ 0, *b*_1_ = *b*_2_ = 0). In case three and four, threats impact both parameters. Case three only considers interactions between parameters ( *a*_1_, *b*_1_ ≠ 0, *a*_2_ = *b*_2_ = 0) while case four considers both interactions between and within both parameters ( *a*_1_, *a*_2_, *b*_1_, *b*_2_ ≠ 0).

Analytical analysis of these equations is difficult to interpret, consequently, we use simulations to further investigate the conditions for additivity through simulations. We simulate 10^6^ random populations (randomly chosen values for r from a uniform distribution between 0 and 1) at different magnitudes (0 to 1) of the impact over 1000 timesteps, to reach the equilibrium population size. Since the harvesting ratio 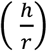 changes the magnitude of the impact at which the additivities occur, we have chosen specific values for the harvesting ratio and split the simulations according to those values, to enable better visualisation. Those values are 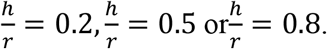 Since *r* is chosen randomly, *h* is assigned to each simulation so that 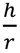 equals the value specified for each group. Equilibria are re-calculated three times for each random population with three different treatments: each one of the threats acting separately, and then the two threats interacting. The additivity index of the two threats for each population is calculated (Eq. 4).

Finally, we simulated the effects of management on all four cases by decreasing the impact of one or both threats by 5% and recalculating the long-term population equilibrium. Management actions could be designed to reduce the threat as a whole, for example reducing fishing pressure, or to reduce the impact on one population parameter, for example fishing technique is changed so that less habitat destruction is caused. For simplicity, it is assumed here that a management action reduces the impact of a threat on both population parameters simultaneously and equally. The benefit is recorded for random populations and across the magnitude of threat impacts of all cases (∼100,000 data points per case)

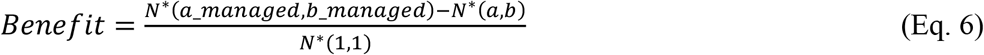

## Results

The results of the simulations agree with the results of the analytical analysis; consequently both are appropriate for analyzing the threat interactions. However, caution has to be given to the defined parameter space to prevent negative population sizes.

Cases 1 and 2 both concentrate on one of the population parameters (Fig 2). These two cases show very different patterns in the joint damages. Two impacts on the carrying capacity always cause a joint damage that is additive until the extinction line (at which point at least one threat can cause extinction) where the joint damage becomes necessarily subadditive. The impacts on the growth rate, however, display a joint damage that is additive at low impacts and superadditive at high impacts. Within the area of extinctions there is subadditive joint damage. Generally, it can be said that the additivity index decreases from zero towards −1 until it hits the extinction line, then the additivity index starts to increase until it reaches +1. As harvest levels increase (Fig 2), the extinction line moves closer towards the origin, as extinction occurs at lower threat impact levels.

**Figure 2.**
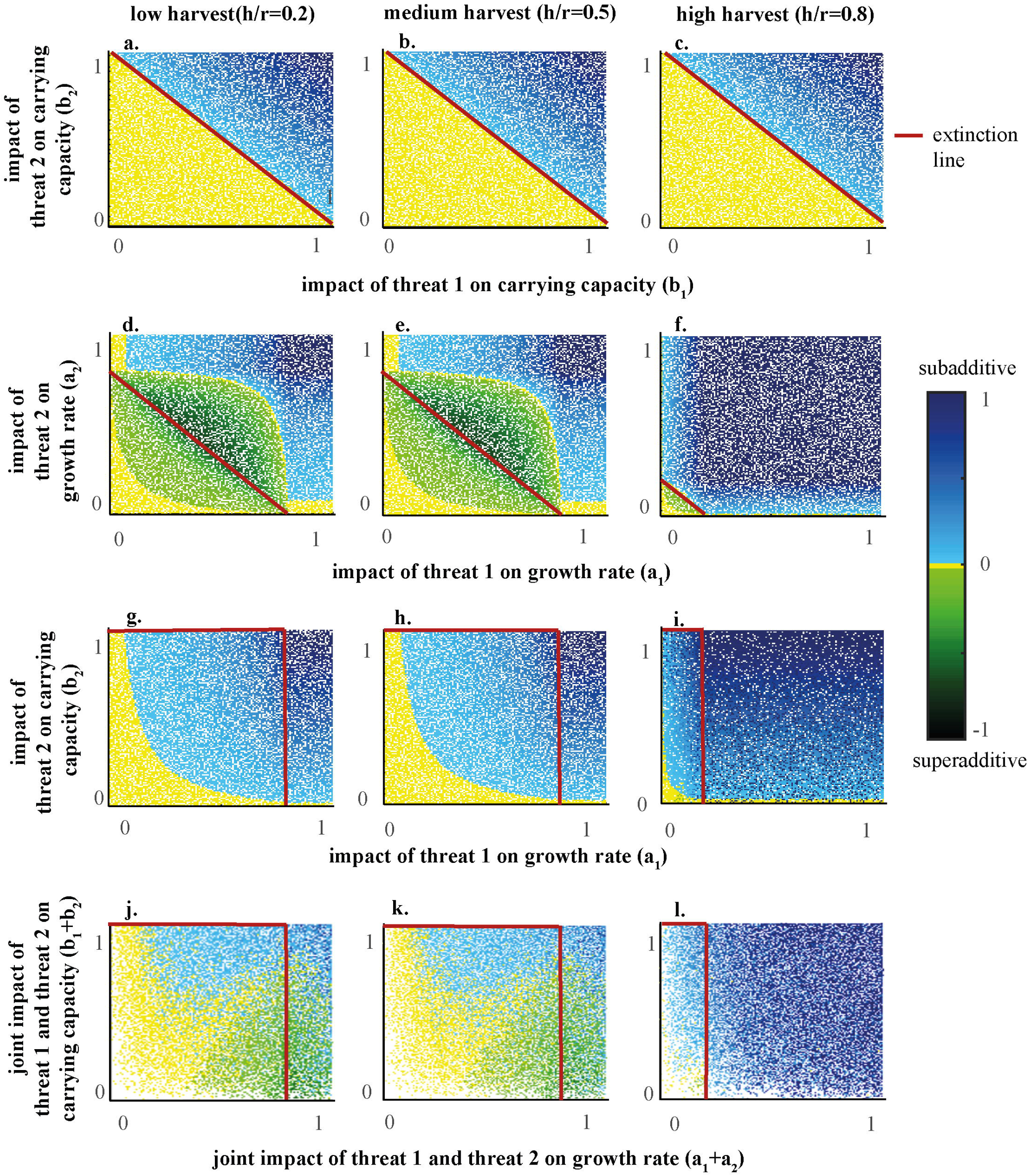
Additivity indices for 10^6^ simulations of random values for h and r split into three cases depending on the parameter impacted by the threats. The four cases represent: a-c: Case 1; Two threats that only impact the carrying capacity ( *a*_1_ = *a*_2_ = 0, *b*_1_, *b*_2_ ≠ 0); d-f: Case 2; Two threats that only impact the growth rate ( *a*_1_, *a*_2_ ≠ 0, *b*_1_ = *b*_2_ = 0); g-i: Case 3; Each parameter is only impacted by one threat ( *a*_1_, *b*_1_ ≠ 0, *a*_2_ = *b*_2_ = 0); j-l: Case 4; Both threats impact both parameters ( *a*_1_, *a*_2_, *b*_1_, *b*_2_ ≠ 0). The columns indicate the level of harvest relative to the population growth rate. Between the origin and the extinction line, the population of organisms persists in the present of the threats, from the extinction line onwards, the population will go extinct in the presence of at least one threat in isolation. The interpretation of an additivity index of zero has to be done carefully, since the graph aligns all values in the range −0. 02 < 0 < 0. 02 as zero.

Case 3 demonstrates the joint damage when both parameter are impacted. This case displays only additive and subadditive damage similar to case 1. However, subadditivity now also occurs without the presence of extinction. Furthermore, it is interesting to note that changes to the impact on the growth rate (*a*_1_) and changes to the impact on the carrying capacity (*b*_1_) do not cause the same change in the additivity index.

In case 4, both threats impact both the carrying capacity and the growth rate. This means that we can compare it directly with case 1 and 2, since they are basically a subset of the simulations displayed within case 4. The only difference is that the results are collapsed into a lower dimensional space. For example, case 1 shows the impact of threat 2 on the carrying capacity on the y-axis and the impact of threat 1 on the carrying capacity on the x-axis. In case 4, both impacts are displayed on the y-axis by simple addition. This means that when the impact on the growth rate is very low in case 4 (*a*_1_ + *a*_2_ = 0), then case 4 is equivalent to case 1. Similarly we can find the results from case 2 in case 4 by setting the impact on the carrying capacity close to zero (*b*_1_+ *b*_2_ = 0). The rest of the case 4 compromises a mixture of sub- and superadditivity. Superadditivity occurs only at high impacts on the growth rate, while subadditivity mainly occurs at medium impacts on both the growth rate and carrying capacity. The absolute magnitude of the additivity index still increases towards the extinction line with increasing impacts of one or either population parameters.

Next, we consider the relationship between the population parameter and the population equilibrium (Fig 3). When increasing the impacts of threats (e.g. from 0 to 1) on the growth rate we can see a decrease in the population equilibrium (Fig 3A). This decrease is first slow then becomes steeper resulting in a concave relationship. On the other hand, when increasing the impacts on the carrying capacity the population equilibrium decreases linearly (Fig 3B). Finally, when we increase the impact on both parameters at different levels (Fig 3C), we can identify all three; slightly convex (red line), piecewise linear so also convex (dark blue line), linear (light blue line) and concave (green line) relationships (Fig 3D).

**Figure 3.**
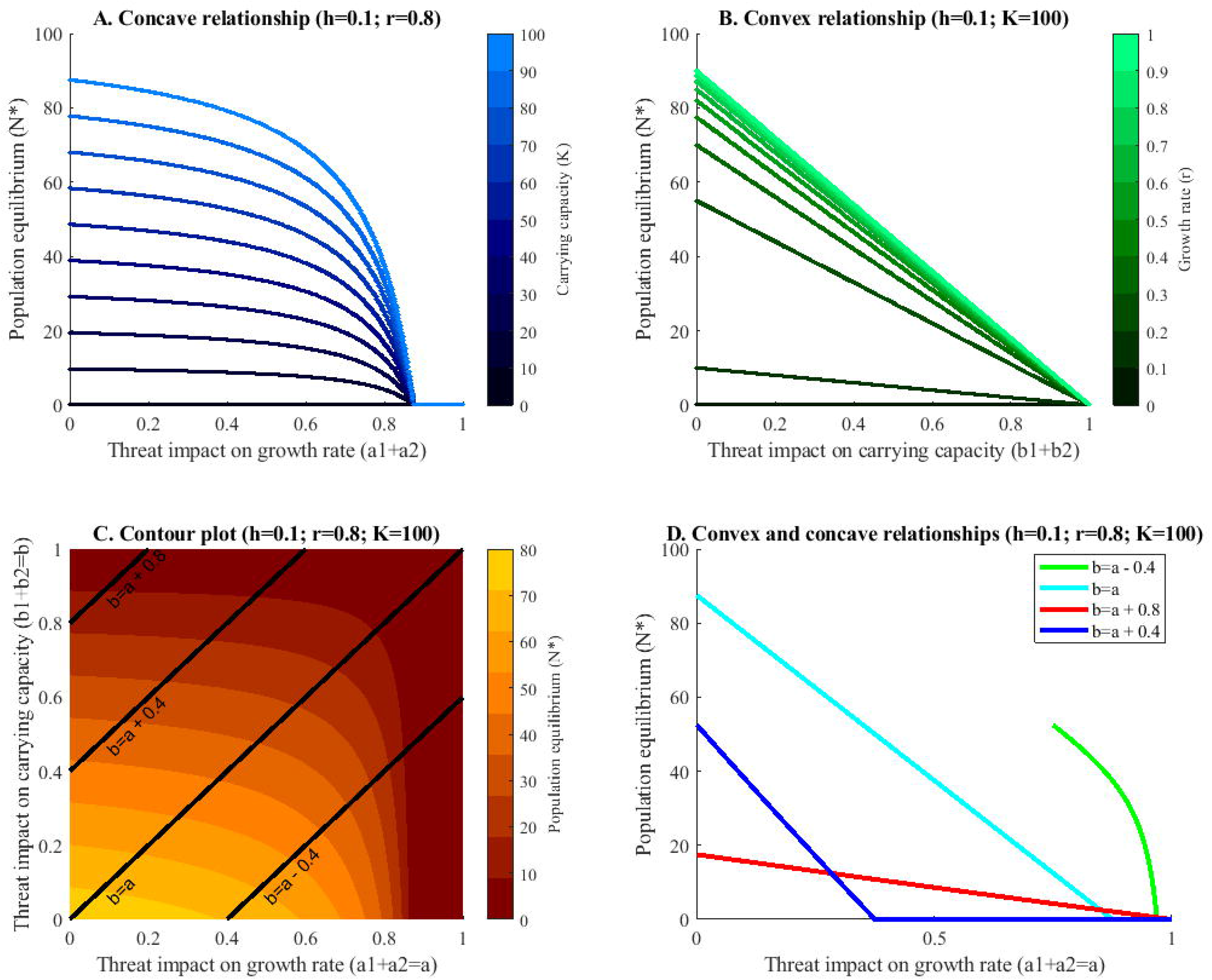
Relationships between the threat impacts on (A) growth rate, (B) carrying capacity, (C&D) growth rate and carrying capacity and the population equilibrium. Part A. shows a concave relationship between the threats impacting the growth rate and the population equilibrium for all levels of the carrying capacity. Part B shows a linear relationship between magnitude of impact on the carrying capacity and the population equilibrium for all magnitudes of the growth rate. Part C shows a contour graph of the population equilibrium with varying threats impacting the growth rate and the carrying capacity. Furthermore, slices are highlighted (lines) that are displayed in Part D. The x-axis in part D represents the magnitude of the impact of the threats on the growth rate. The threats impacting the carrying capacity are also varied and can be identified using the appropriate linear function to calculate b. Part D shows that depending on the slice we choose from Part C both concave and convex (here piecewise linear) relationships can be found when all threat impacts are varied.

Management benefit per 5% impact change displays a large variation from as low as 0% increase of the no threat population equilibrium up to 4300% increase (Table 3). Both extremes occur when the equilibrium populations are close to zero before management.

**Table 3.**
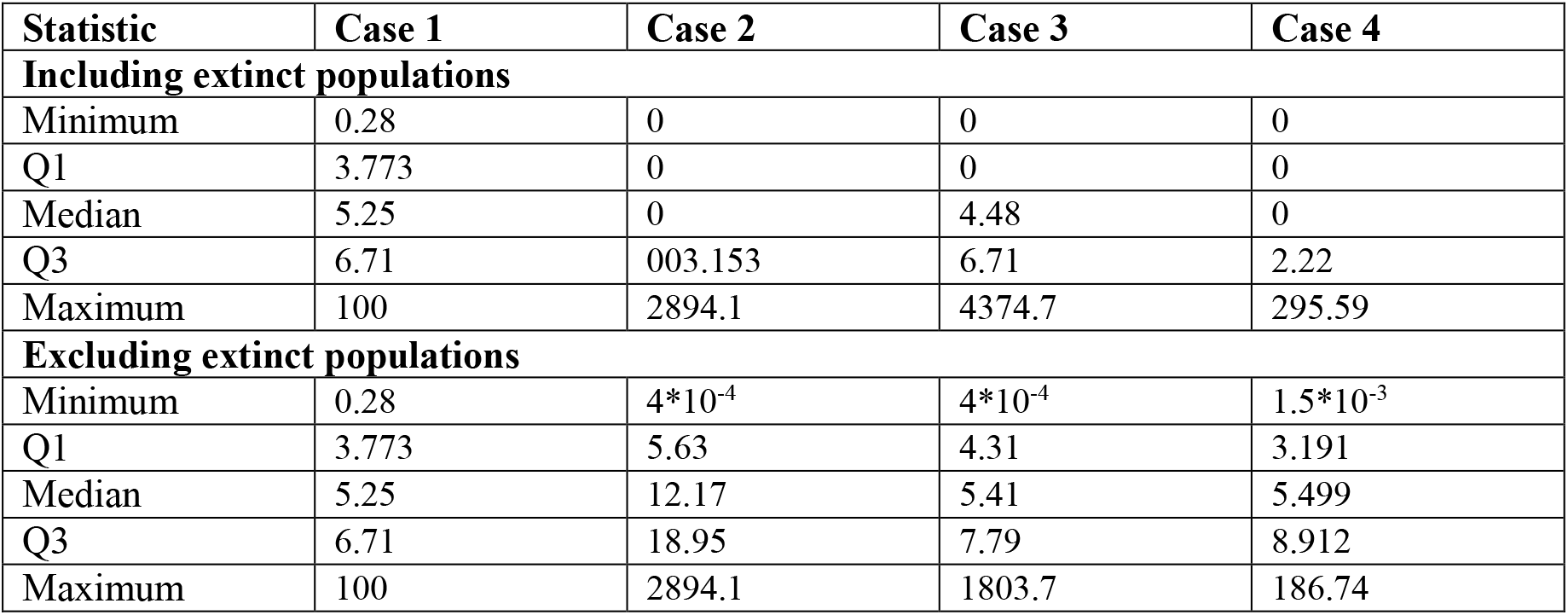
Statistics summarising all of the simulations used for Fig. 4 divided according to the cases and extinction status after the management on the threats.

Additivity of multiple threats
| Statistic | Case 1 | Case 2 | Case 3 | Case 4 |
| --- | --- | --- | --- | --- |
| **Including extinct populations** |  |  |  |  |
| Minimum | 0.28 | 0 | 0 | 0 |
| Q1 | 3.773 | 0 | 0 | 0 |
| Median | 5.25 | 0 | 4.48 | 0 |
| Q3 | 6.71 | 003.153 | 6.71 | 2.22 |
| Maximum | 100 | 2894.1 | 4374.7 | 295.59 |
| **Excluding extinct populations** |  |  |  |  |
| Minimum | 0.28 | $4 \cdot 10^{-4}$ | $4 \cdot 10^{-4}$ | $1.5 \cdot 10^{-3}$ |
| Q1 | 3.773 | 5.63 | 4.31 | 3.191 |
| Median | 5.25 | 12.17 | 5.41 | 5.499 |
| Q3 | 6.71 | 18.95 | 7.79 | 8.912 |
| Maximum | 100 | 2894.1 | 1803.7 | 186.74 |

Several factors influence the management benefit experienced by a population when particular threats are decreased. First, there is the magnitude of the impact. The impact on the parameter growth rate shows some variation with benefit being higher in the extreme case (high and low magnitude) versus the medium magnitude (Fig 4a). On the other hand, the impact on the carrying capacity shows a clear decrease of management benefit with a decrease in magnitude. The lowest impact doubles or even triplets the management benefit experienced (Fig 4b). The largest amount of variation is explained when we consider the additivity together with the benefit (Fig 4c). More superadditive behavior lead to over ten times the benefit compared to cases where very subadditive damage is displayed.

**Figure 4.**
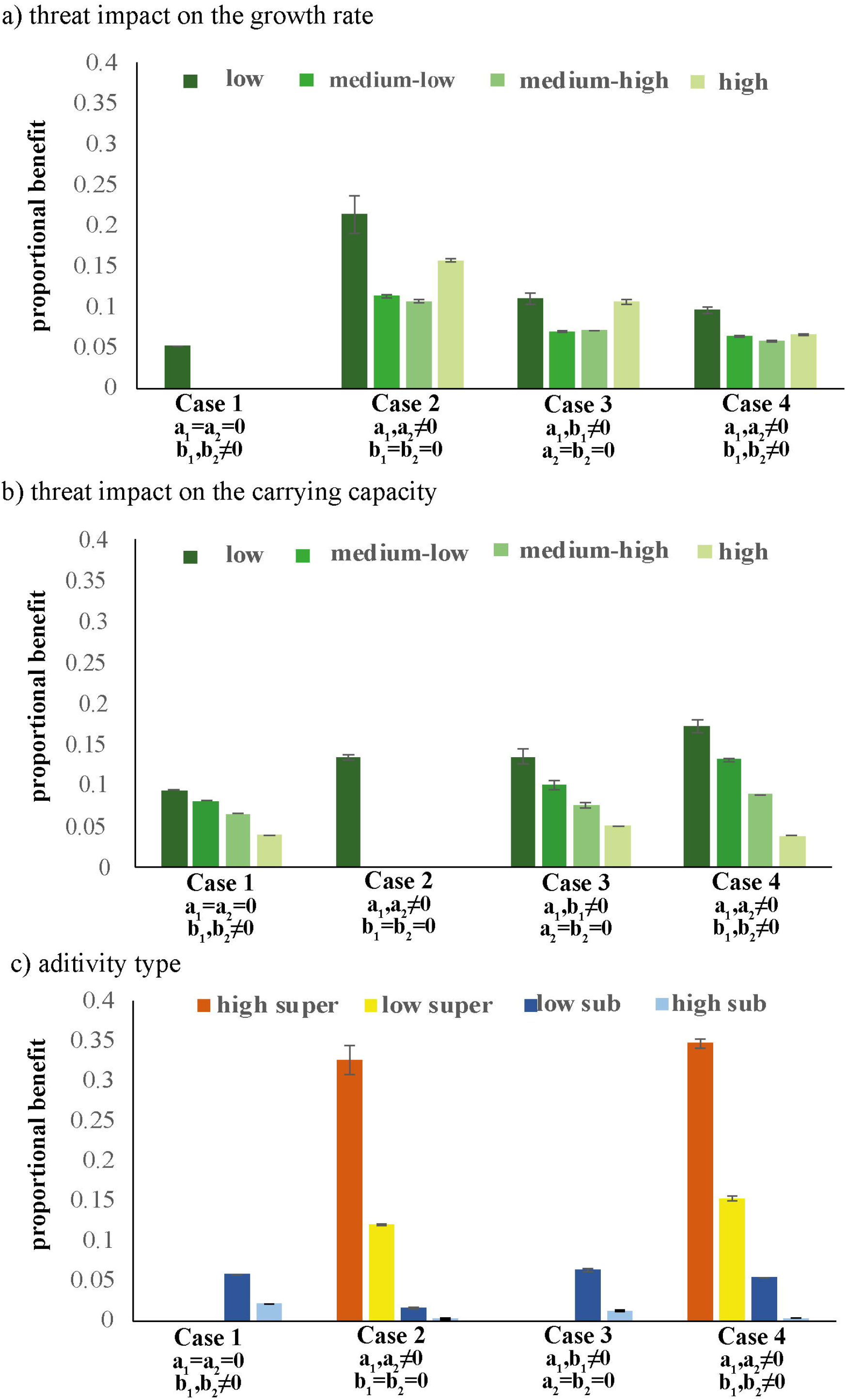
Management benefit (±1.96*SE) when reducing both threat impacts simultaneously according to the four cases. The results for the cases are split on the x-axis. Panel a) splits the benefit for different magnitudes of threat impact on the parameter carrying capacity. Panel b) splits the benefit for different magnitudes of threat impacts on the growth rate. Low impact < 0.25; 0.25< Medium-Low impact < 0.5; 0.5 < Medium-High impact < 0.75; High impact > 0.75. Panel c) splits the benefit depending on the additivity type. High superadditivity < −0.5; −0.5 < Low superadditivity < 0; 0 < Low subadditivity < 0.5; High subadditivity > 0.5.

## Discussion

This study explored the interaction behavior of two threats acting upon two population parameters in theoretical populations. We found that, contrary to orthodox assumptions, the additive dynamics are not inherent to the particular threat combination (7, 9). Even in a simple, one-species model, the additivity can exhibit qualitative changes, depending on the affected parameter, and the magnitude of the impact on a threat. Our results therefore suggest that studies or reviews should be careful when they attribute the qualitative type of additivity to particular combinations of threats (4), and be aware that the parameters affected and the magnitude of the impact could be driving the additive dynamics

In our models, superadditivity only occurs if there are several impacts on the growth rate. This can be explained by the concave relationship between the intrinsic growth rate and the equilibrium population size (Fig. 3a). Following this curve toward the origin, we see that the slope increases as the magnitude of the impact increases. A threat with twice the impact will therefore cause more than double the damage to the equilibrium population size. In contrast, the joint damage of threats will be additive when the slope is constant, i.e. a linear relationship between the population parameter and the population equilibrium (Fig. 3b). When we consider the multiplicative model of impacts, this relationship becomes convex indicating subadditivity (see supplementary materials S1).

During this study, we found a few occasions where concavity does not predict all of the interactions that we can find, i.e. this generalization seems to contradict our results. For example, at high magnitude of impacts there are subadditive interactions. On closer examination, we found that all of these subadditive data points were associated with extinctions. This subadditivity can only be found in the simulation results, the analytical analysis results in negative population sizes, which are not ecologically defined. Consequently, the data points resulting in extinction (negative population sizes, subadditivity) lay outside the realm of definition of the concave function.

Interestingly, reducing both parameters simultaneously can cause both super- and subadditivity at varying magnitudes of threat impacts. This is also reflected in the parameter-equilibrium relationships that can be both convex and concave (Fig. 3c-d). This means that at high levels of the growth rate and the carrying capacity the curve is concave, causing superadditivity and at low levels convex, causing subadditivity without extinction.

This confirms our results and leads to the conclusion that we can infer the additive behavior from the curvature of the applicable curve.

It is interesting to note that we found more differences in additivity of damages due to the magnitude and parameters affected by impacts rather than defining the impacts as additive or multiplicative. This highlights the importance of differentiating between impacts and damages in any studies concerning threats.

Additivity of multiple threats has been considered in terms of conservation and management of populations repeatedly. In many cases, the opinion is that superadditivity is the worst case for the population (15, 16). However, superadditivity can also be the best case scenario when considered from the perspective of management (17). Our results support this since threats with measured superadditivity result in the largest proportional management benefit. This can be especially true when we consider local versus global, manageable versus unmanageable, threats. Superadditivity can mean that by reducing a manageable threat we can simultaneously achieve a reduction in the damage caused by the unmanageable threat (5). On the other hand subadditivity would mitigate the benefit from the management of a single threat and, consequently, the management action would be of less use than an equivalent superadditive situation. Generally speaking the management benefits are easiest to predict for an additive threat combination (9), however, if we know and respect the underlying additive dynamics as described in this paper, predictions for other situations can become more accurate.

These results in combination with the commonly-conducted cumulative threat mapping (6, 18) can be used to prioritise management actions. Prioritising management is especially important in ecosystems that spread over large areas where it is impossible to protect the full extent of a species (19). In such systems prioritising management actions is crucial. When prioritising there are many aspects to consider, such as cost, risk, suitability and resulting benefit (20). The analysis shown here can aid in the assessment of the suitability for management of different areas and likely benefit that can be achieved. Global threats are always difficult to manage for local government so are less suitable. So if a global threat impacts all areas of conservation concern, but different local threats impact specific areas, then according to the analysis here, we might want to protect the areas that impact in a fashion (magnitude and parameter impacted) that cause superadditivity (all else being equal). Furthermore, the actual benefit that a management action can result in is influenced by all threats to this system. The analysis conducted here, i.e. knowledge of the parameters impacted by each threat can help to estimate likely benefits. Therefore, these findings could streamline some aspects of management prioritisations.

As with any theoretical analysis, this study is based on a number of simplifications. Most importantly, it utilises a simple logistic model that considers one population is isolation. This is not particularly realistic since species interactions are ubiquitous and important, and threats can also interact with each other through those species. However, for this study a simple model is used to highlight the complexities that interactions introduce. It is important to note here that a more complex model will result in more complexities in the result not less. The simple model also provides a framework to interpret and explain some of the phenomena that are likely to still play role in more complex communities. The applicability of these results for many populations is also confirmed through the use of the Beverton-Holt and the Ricker model that both showed the same patterns of additivity (S1). Future work will aim to transfer the conclusions and explanations from a single population in this study to more complex community level models (connection between curvature and additivity in damages).

Second, our calculations are based on population equilibria when exposed to different combinations and magnitudes of impacts. Population equilibria are used regularly in many ecological models, but are also countered by many (20–23). The main criticism argues about the oscillations around populations equilibria. While these are justified, it is often commented that the usefulness of equilibria as a base assumption depends on the scale (24, 25). At small spatial scales the dynamics are more transient, while at large spatial scales dynamics stabilise to render the equilibrium assumption realistic enough (26). Used appropriately, investigations of the equilibrium can provide useful insights.

## Conclusions

This study has provided an overview of the complexity of behaviors that interacting threats can display. Overall, the traditional idea of assigning different additive dynamics to particular threat combinations, is expanded to also include the parameters impacted, and the magnitude of those impacts. Besides the complexity revealed in this study, insights can be drawn about the origins of superadditive behavior: in our models, this important dynamic only occurs when multiple threats impact the growth rate of a population. More generally, the interaction behavior can be predicted by the curvature of the relationship between the impacted parameter and the equilibrium population size; a convex relationship implies subadditivity, and a concave relationship implies superadditivity. Finally, this study urges ecologists to focus on identifying the parameter and relative magnitude of impacts for each threat combination rather than just the additivity type.

## SUPPORTING INFORMATION

**S1 Multiplicative impact on the logistic model**

**S2 Additive and multiplicative impact on the Beverton-Holt model**

**S3 Additive and multiplicative impact on the Ricker model**

