## Supplementary material for "Superadditive and subadditive dynamics are not inherent to the types of interacting threat": S1

### S1 Multiplicative impact on the logistic model

#### Justification

In the main manuscript we describe that impacts can be modelled as either multiplicative or additive. Which one is more suitable might depend on the actual threat or the physiology of the species. The purpose of this supporting material is to show that the results and conclusions do not depend on the assumptions of additive impacts made in the main paper.

#### Methods

The logistic model used remains the same, i.e. the model used is

$$\frac{dN}{dt} = rN\left(1 - \frac{N}{K}\right) - hN \qquad \text{(S1-1)}$$

Consequently, the population equilibrium in the presence of a threat is

$$N^*(a, b) = \left(1 - \frac{h}{(1-a)r}\right)(1-b)K. \qquad \text{(S1-2)}$$

To reflect the multiplicative nature of the impact they are now modeled as following

$(1 - a_1) * (1 - a_2)$, and $(1 - b_1) * (1 - b_2)$.

#### Results

Some differences can be seen between Figure 2 in the main paper and Figure S1-1. The main one here is that the additivity index shows more positive values (i.e. subadditivity) than zero values. This is caused by the multiplication of threat impact, i.e. the subadditivity introduced within the impact. This can also be seen in Figure S1-2B with a convex curvature that is associated with subadditivity.

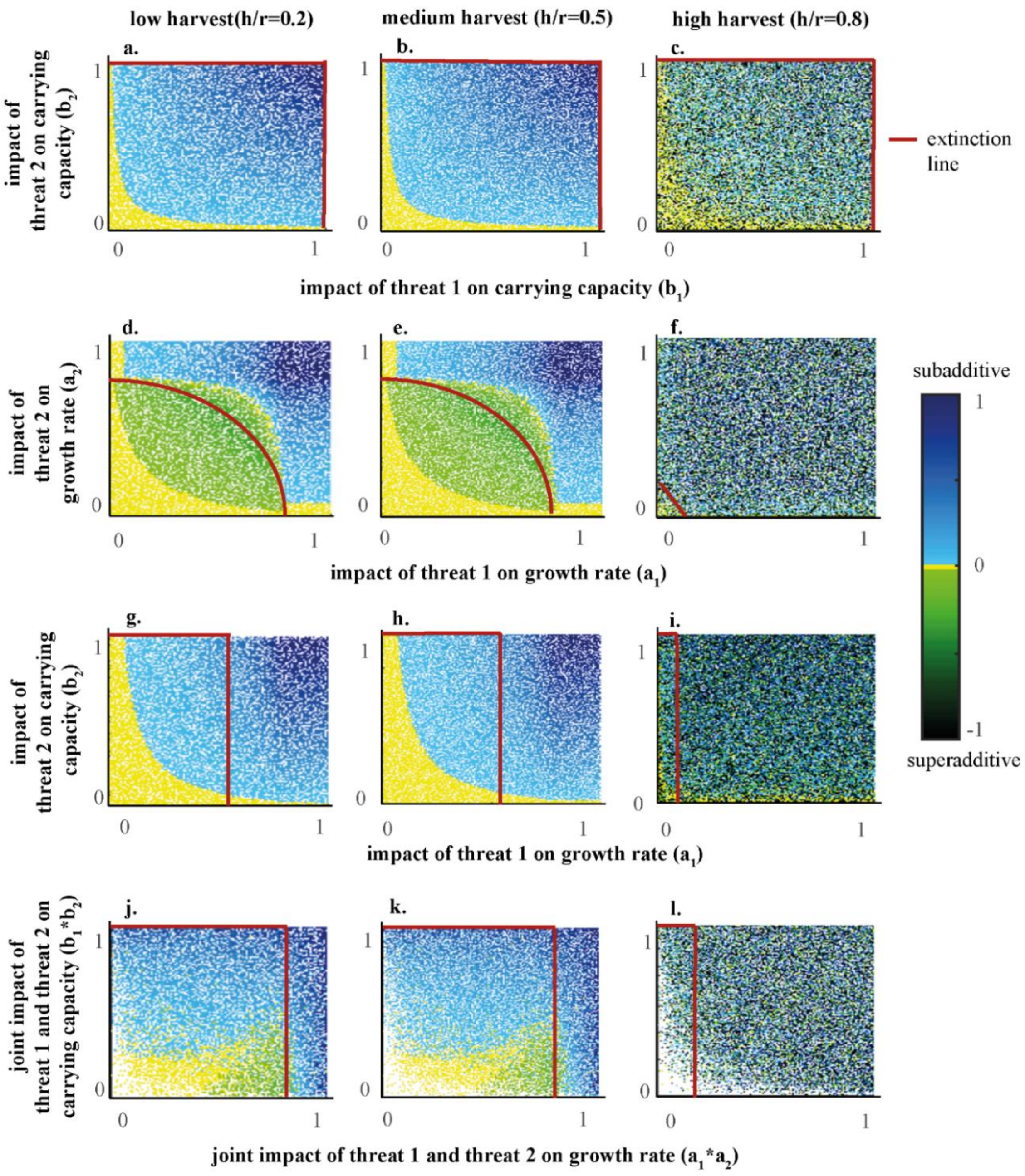


**Figure S1-1. Additivity indices for $10^6$ simulations of random values for h and r split into three cases depending on the parameter impacted by the threats. The four cases represent: a-c: Case 1; Two threats that only impact the carrying capacity ( $a_1 = a_2 = 0, b_1, b_2 \neq 0$ ); d-f: Case 2; Two threats that only impact the growth rate ( $a_1, a_2 \neq 0, b_1 = b_2 = 0$); g-i: Case 3; Each parameter is only impacted by one threat ( $a_1, b_1 \neq 0, a_2 = b_2 = 0$) ; j-l: Case 4; Both threats impact both parameters( $a_1, a_2, b_1, b_2 \neq 0$). The columns indicate the level of harvest relative to the population growth rate. Between the origin and the extinction line, the population of organisms persists in the present of the threats, from the extinction line onwards, the population will go extinct in the presence of at least one threat in isolation. The interpretation of an additivity index of zero has to be done carefully, since the graph aligns all values in the range $-0.02 < 0 < 0.02$ as zero.**

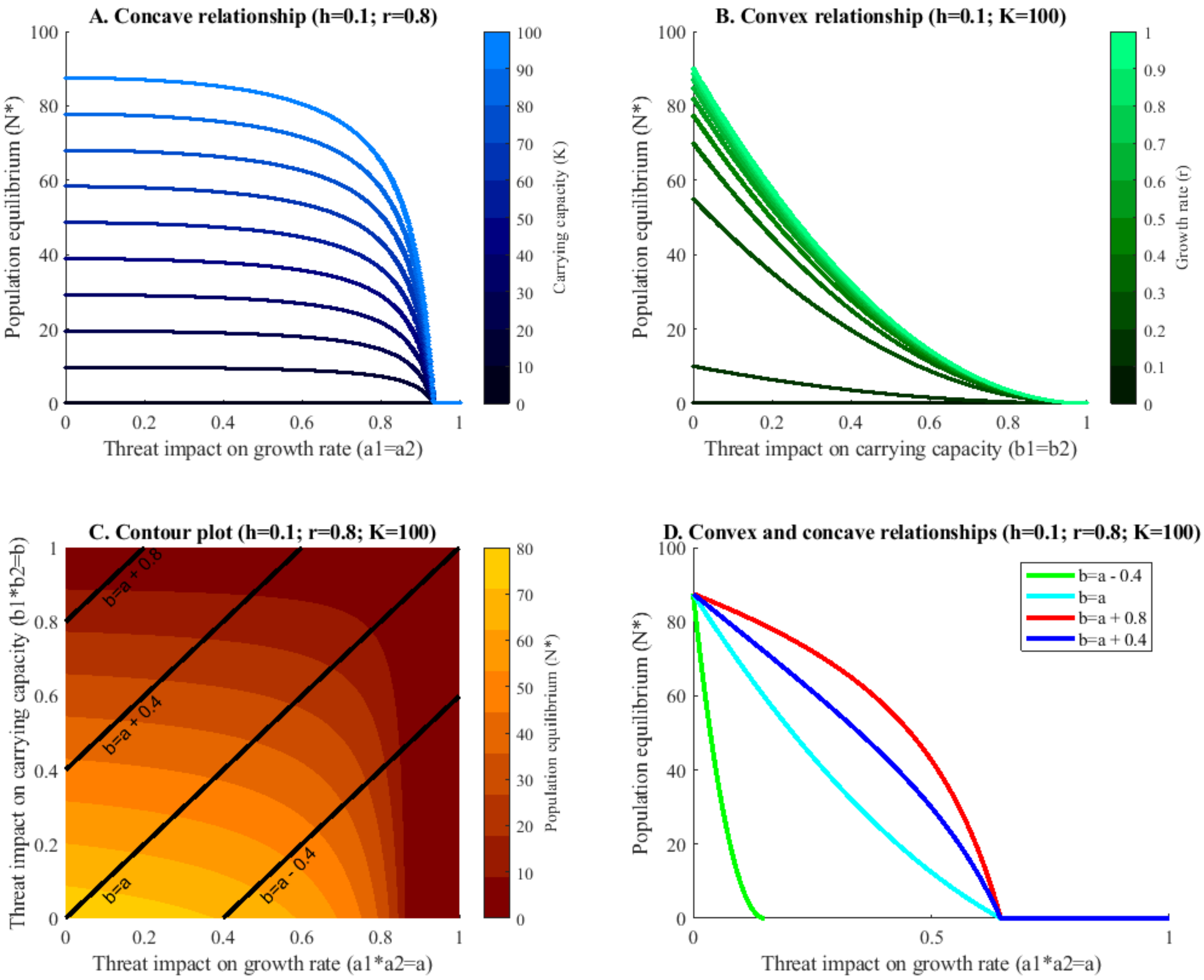


**Figure S1-2. Relationships between the threat impacts on (A) growth rate, (B) carrying capacity, (C&D) growth rate and carrying capacity and the population equilibrium. Part A. shows a concave relationship between the threats impacting the growth rate and the population equilibrium for all levels of the carrying capacity. Part B shows a linear relationship between magnitude of impact on the carrying capacity and the population equilibrium for all magnitudes of the growth rate. Part C shows a contour graph of the population equilibrium with varying threats impacting the growth rate and the carrying capacity. Furthermore, slices are highlighted (lines) that are displayed in Part D. The x-axis in part D represents the magnitude of the impact of the threats on the growth rate. The threats impacting the carrying capacity are also varied and can be identified using the appropriate linear function to calculate b. Part D shows that depending on the slice we choose from Part C both concave and convex (here piecewise linear) relationships can be found when all threat impacts are varied.**
