## Supplementary material for "Superadditive and subadditive dynamics are not inherent to the types of interacting threat": S2

### S2 Additive and multiplicative impact on the Beverton-Holt model

#### Justification

In the main manuscript we use the logistic model. However, other models can be used to model single species. One of them is the Beverton-Holt model first proposed in 1957. The aim of this supplementary material is to show that the conclusions in the main text hold for the Beverton-Holt model.

#### Methods

The Beverton-Holt model is a discrete time model given by

$$N_{t+1} = \frac{N_t(r-1)}{1+\frac{N_t r}{K}} - hN_t \qquad \text{(S2-1)}$$

with the growth rate ($r$), the harvest rate ($r$) and the carrying capacity ($K$). This specific construction of the Beverton-Holt model aassigns harvest to occur before growth.

Consequently, the population equilibrium in the presence of a threat is

$$N^*(a,b) = [(1-h)((1-a)r+1)-1]\frac{(1-b)K}{(1-a)r}. \qquad \text{(S2-2)}$$

Here we present both the additive nature of the impact, $(1-a_1)+(1-a_2)$ and $(1-b_1)+(1-b_2)$,as well as the multiplicative nature of the impact $(1-a_1)*(1-a_2)$, and $(1-b_1)*(1-b_2)$.

#### Results

Both of the figures here are barely differentiable results to the logistic model. So the conclusions hold for the Beverton-Holt model as well.

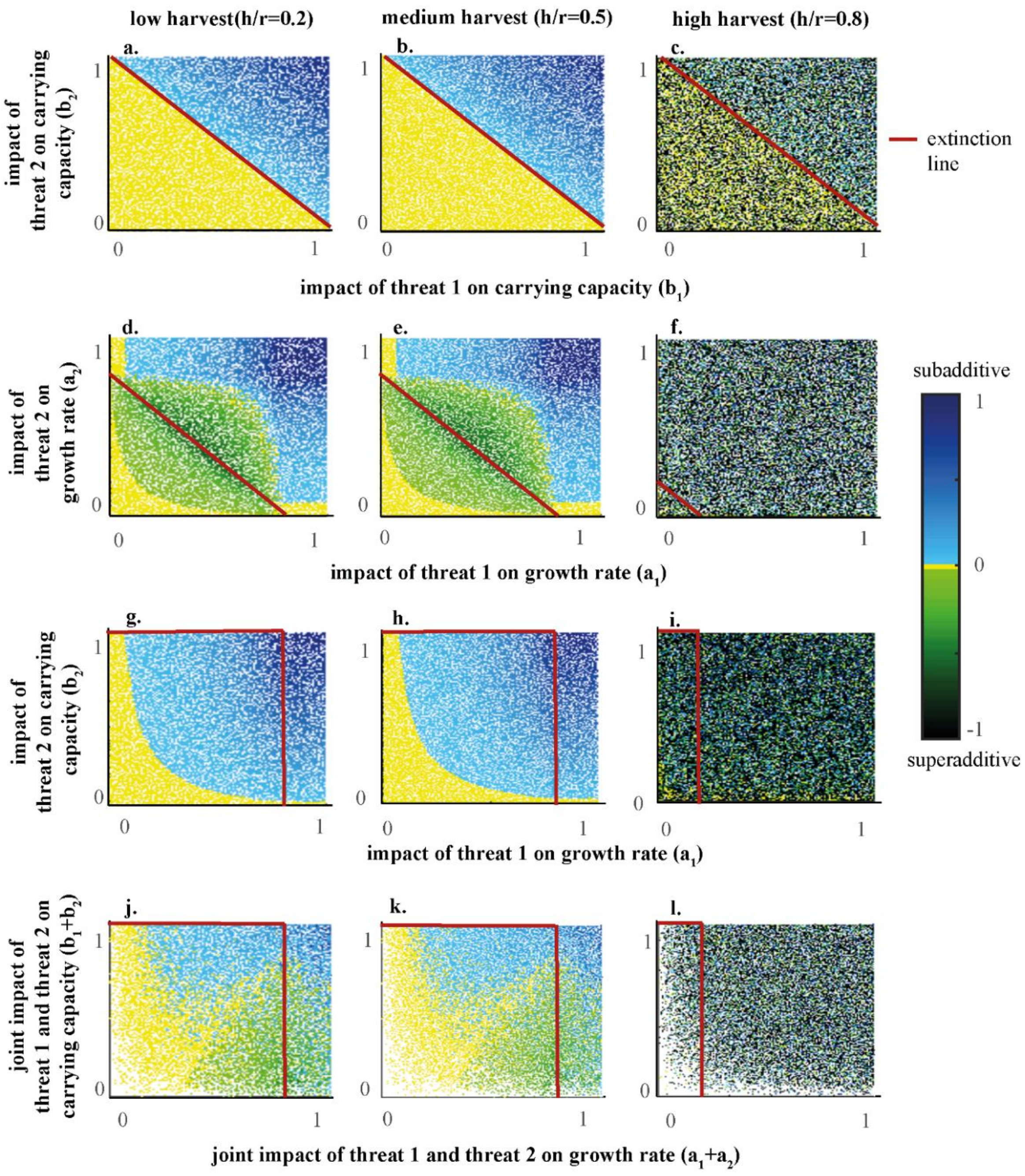


**Figure S2-1. Additivity indices for $10^6$ simulations of random values for h and r split into three cases depending on the parameter impacted by the threats. The four cases represent: a-c: Case 1; Two threats that only impact the carrying capacity ( $a_1 = a_2 = 0, b_1, b_2 \neq 0$ ); d-f: Case 2; Two threats that only impact the growth rate ( $a_1, a_2 \neq 0, b_1 = b_2 = 0$); g-i: Case 3; Each parameter is only impacted by one threat ( $a_1, b_1 \neq 0, a_2 = b_2 = 0$) ; j-l: Case 4; Both threats impact both parameters( $a_1, a_2, b_1, b_2 \neq 0$). The columns indicate the level of harvest relative to the population growth rate. Between the origin and the extinction line, the population of organisms persists in the present of the threats, from the extinction line onwards, the population will go extinct in the presence of at least one threat in isolation. The interpretation of an additivity index of zero has to be done carefully, since the graph aligns all values in the range $-0.02 < 0 < 0.02$ as zero.**

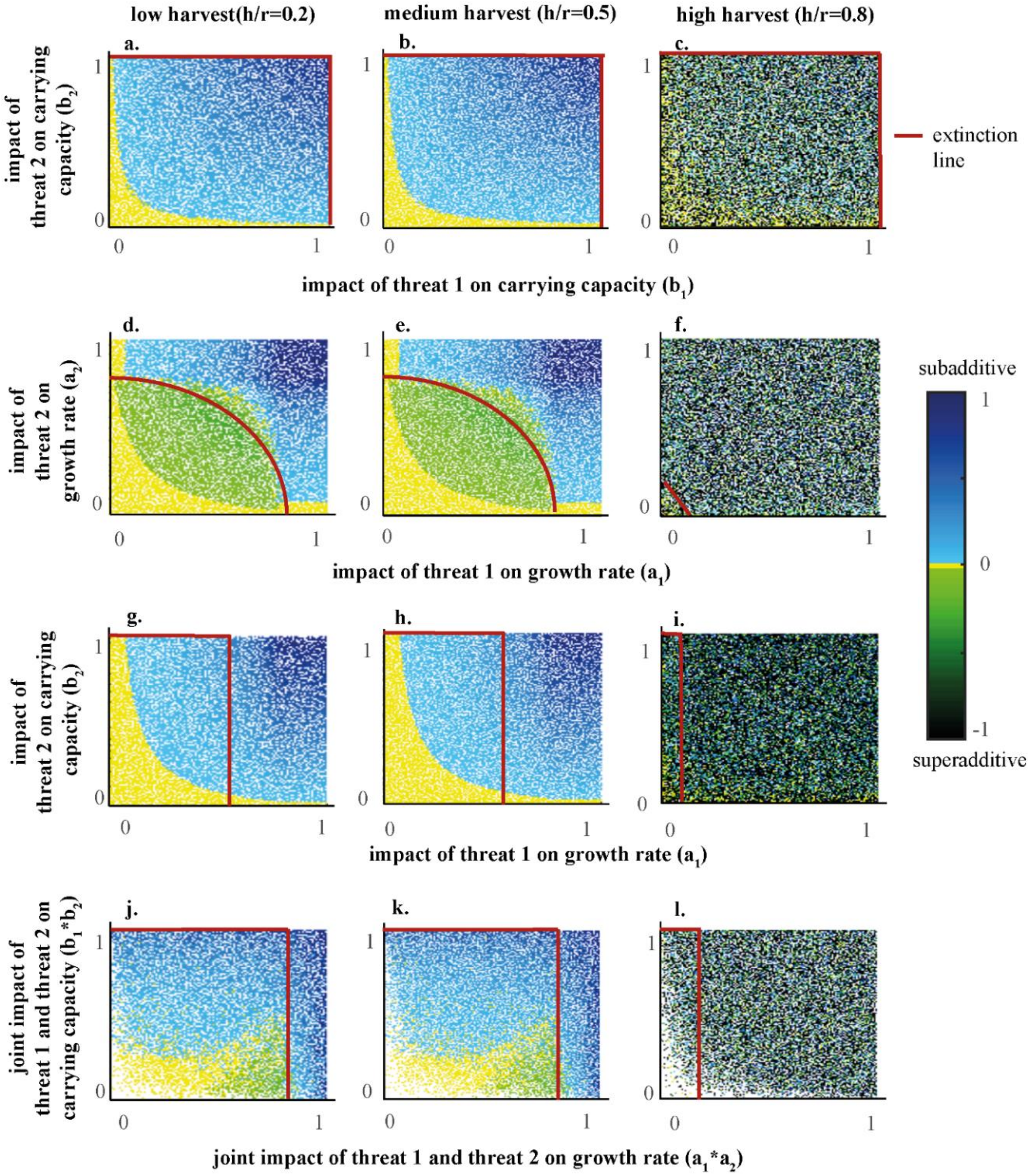


**Figure S2-2. Additivity indices for $10^6$ simulations of random values for h and r split into three cases depending on the parameter impacted by the threats. The four cases represent: a-c: Case 1; Two threats that only impact the carrying capacity ( $a_1 = a_2 = 0, b_1, b_2 \neq 0$ ); d-f: Case 2; Two threats that only impact the growth rate ( $a_1, a_2 \neq 0, b_1 = b_2 = 0$); g-i: Case 3; Each parameter is only impacted by one threat ( $a_1, b_1 \neq 0, a_2 = b_2 = 0$) ; j-l: Case 4; Both threats impact both parameters( $a_1, a_2, b_1, b_2 \neq 0$). The columns indicate the level of harvest relative to the population growth rate. Between the origin and the extinction line, the population of organisms persists in the present of the threats, from the extinction line onwards, the population will go extinct in the presence of at least one threat in isolation. The interpretation of an additivity index of zero has to be done carefully, since the graph aligns all values in the range $-0.02 < 0 < 0.02$ as zero.**
